# New insight into erythema reduction induced by quercetin metal ions chelates

**DOI:** 10.1101/2024.12.04.626931

**Authors:** Alain Bolaño Alvarez, Kristian B. Arvesen, Kasper F. Hjuler, Peter Bjerring, Steffen B. Petersen

## Abstract

In dermatology, chelates play a significant role in skin care and treatment of skin diseases. Chelates involve coordination bonding between metal ions and organic molecules such as flavonoids. Quercetin (**Q**) is an extensively studied natural flavonoid with proven safety and anti-inflammatory properties. This study shows the role of two biocompatible metal ions, Iron (**Fe**^**2+**^) and Copper (**Cu**^**2+**^) in coordination bonding with **Q** in 2-Propanol 50% and 80% at 1:1 stoichiometry. Our results show that chelation involves the hydroxyl groups and occurs by coordination of **Cu**^**2+**^ to **Q** for the Ring A-B (Benzoyl group) resulting in a fluorescence emission peak at 530nm from the Ring B-C (Cinnamoyl group). This chelates **Q+Cu**^**2+**^ reduces mechanically induced erythema in the skin (Tanned type). A similar effect was observed in the chelate **Q+Fe**^**2+**^ where the coordination of **Fe**^**2+**^ to **Q** occurs for the cinnamoyl group resulting in an emission peak at 425nm from Benzoyl group of **Q**. The statistical analysis shows significant differences in the effects of **Q+Cu**^**2+**^ (p-value = 0.00001), **Q+Fe**^**2+**^ (p-value = 0.0003) respect to **Q** as well as between them (p-value =0.0029). Our results suggest that the interaction between **Q** and metal ions plays a central role in the inflammatory pathway. We conclude that the anti-inflammatory properties of **Q** were enhanced by both **Q+Fe**^**2+**^ and **Q+Cu**^**2+**^ chelates, highlighting the effect of **Q+Fe**^**2+**^ where the hydroxyl groups available in the cinnamoyl group of the **Q** molecule are the main intermediates to interact with **Fe**^**2+**^, which is a requirement to trigger the anti-inflammatory molecular events of **Q** molecules.

## INTRODUCTION

Quercetin is a natural molecule (flavonoid) found in various fruits, vegetables, and plants [1, 2]. This molecule can function as a chelating agent owing to the presence of hydroxyl groups in its structure, which can bind to metal ions, particularly transition metals such as iron and copper [3], which are considered biocompatible owing to their physiological effects [4], but can be involved in oxidative reactions [5]. Thus, by chelating metal ions, quercetin can prevent free radical formation, which contributes to oxidative stress and DNA damage, particularly in the case of copper coordination [6]. In the context of dermatology (skin treatment), the chelating capacity of quercetin can have several benefits, such as antioxidant effects [7, 8], anti-aging effect [9] and anti-inflammatory effects [4, 10, 11]. A new investigation that involves chelation offers a window to potential applications in dermatology and a better understanding of the mechanism of chelate action. These metal ions (**Cu**^**2+**^ and **Fe**^**2+**^) have been reported to play distinctive roles in skin health. **Cu**^**2+**^ ions are known for their physiological importance in skin treatments. For example, **Cu**^**2+**^ has been associated with dermal fibroblast proliferation [12], acting as a cofactor for skin enzymes, such as superoxide dismutase [13, 14], and contributing to skin pigmentation through its role in tyrosinase activity, an enzyme important in melanin biosynthesis [15, 16].

In contrast, the role of **Fe**^**2+**^ ions in skin health has been less studied, primarily involving collagen metabolism and procollagen proline dioxygenase [17]. The anti-inflammatory effects of **Fe**^**2+**^ on skin health are less clear than the well-documented iron-chelating effects [18], however, more information is required to better understand the role of **Q** in iron metabolism [19]. However, the iron sequestration capacity of **Q** has been suggested by its photoprotective effects on skin [20]. However, recent studies showed that **Q+Fe**^**2+**^ inhibited inflammatory macrophages in the joints of mice with Rheumatoid Arthritis, reducing the activation of NF-kB pathways [21].

The interactions of **Cu**^**2+**^ and **Fe**^**2+**^ ions with **Q** have also been studied. **Cu**^**2+**^ interacts strongly and preferentially by coordination with the carbonyl oxygen atom, -OH3, and -OH5 groups in the benzoyl group (Ring A-C) of the **Q** molecule [22, 23], but there is evidence of coordination for the cinnamoyl group [3, 24] which depends on the pH of the solution and influences the protonation of the hydroxyl groups [25]. However, the coordination between **Fe**^**2+**^ and **Q** can also occur by the benzoyl group [26], although -OH4’ and -OH3’ in the cinnamoyl group (Ring B) of the **Q** molecule can occur [27-29]. In addition, the strength of the interaction is associated with the pH of the solution, charge of the ions, capacity of the flavonoid to reduce ions [27] and polarity of the solvent [3]. Despite these similarities, **Fe**^**2+**^ has a better capacity to form chelates than **Cu**^**2+**^ and plays an important role in the decomplexation of chelated **Cu**^**2+**^ complexes [30].

In this work, we studied the effects of biocompatible metal ions (**Cu**^**2+**^ and **Fe**^**2+**^) on the spectroscopic properties of **Q**, highlighting the potential of this flavonoid in combination with metal ions to reduce erythema (induced mechanically using a piling tape), a common skin condition characterized by redness and inflammation. In addition, we are driving new insights into quercetin combined with biocompatible metal ions, emphasizing its potential applications in skin health and inflammation reduction by improving the anti-inflammatory properties of **Q**. In addition, we demonstrated that it is possible to distinguish which part of the **Q** molecule (Benzoyl or Cinnamoyl group) is more relevant to trigger the anti-inflammatory effect on the skin, supporting an improved effect of **Q** combined with metal ions.

## RESULTS

### Absorbance properties of quercetin mixtures with Copper and Iron in 50% and 80% 2-Propanol

Is well known that Quercetin (**Q**) is a polyphenol molecule with two characteristic absorption peaks, corresponding to Ring A and Ring B, respectively [31-33]. We examined the absorbance spectra of **Q** and its mixtures with two metal ions, copper (**Q+Cu**^**2+**^), iron (**Q+Fe**^**2+**^), and both copper and iron (**Q+Cu**^**2+**^**+Fe**^**2+**^) in 2-Propanol 50% and 2-Propanol 80%.

In both 2-Propanol 50% and 2-Propanol 80%, the absorbance spectra of Q exhibited distinct peaks at 255nm and 375nm, corresponding to Ring A and Ring B. No significant differences in the intensity of these peaks were observed between 2-Propanol 50% and 2-Propanol 80% (**Figure 1**). However, in **Q+Cu2+** dissolved in both 2-Propanol 50% and 2-Propanol 80%, the absorbance spectrum showed a decrease in the intensity at 375nm (Ring B) compared to Q (**Figure 1A**).

**Figure 1:**
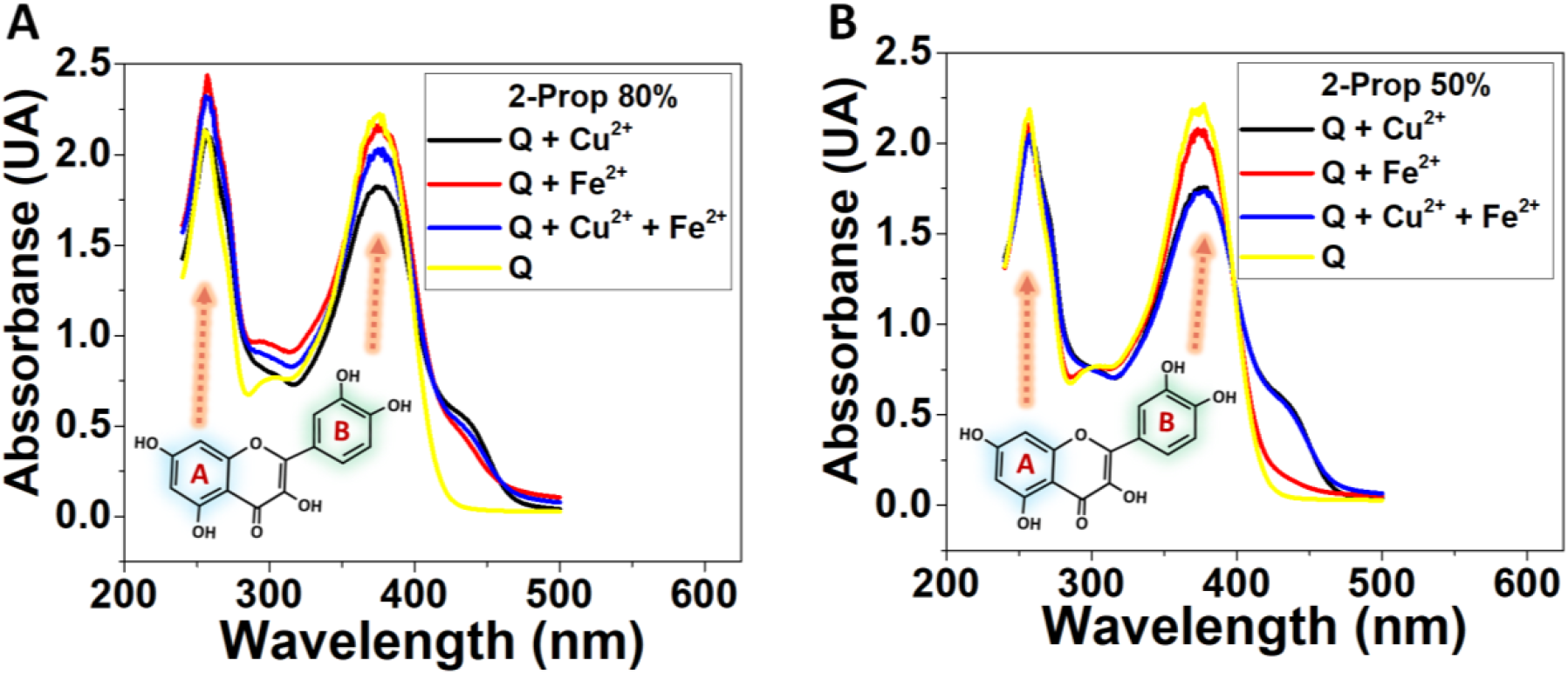
Absorbance properties of Q/Cu^2+^/Fe^2+^ in 2-Prop 80% and 2-Prop 50%. In Q the absorbance peak of Ring A appears at 255 nm and Ring B at 375 nm. Absorbance of the mixtures Q/Cu^2+^, Q/Fe^2+^ and Q/Cu^2+^/Fe^2+^ at 1μM molar proportion each one in 2-Propanol 80% (**A**) and in 2-Propanol 50% (**B**).

Particularly, in **Q+Fe2+** dissolved in 2-Propanol 50% the spectra behave similarly to pure Q without change in the intensity at 375nm (**Figure 1B**). However, in 2-Propanol 80%, the absorbance spectrum of **Q+Fe2+** not showed significant changes in the intensity at 375nm (**Figure 1A**).

In **Q+Cu2+-Fe**^**2+**^ dissolved in both 2-Propanol 50% and 2-Propanol 80% the absorbance spectrum displayed differences compared to pure quercetin (Q), with a decrease in intensity at 375nm (Ring B).

### Thermal Scan measurement of quercetin mixtures with Copper and Iron in 50% and 80% 2-Propanol

The stability of the **Q+Cu2+** and **Q+Fe2+** complexes in 2-Propanol was investigated by a thermal scan from 25°C to 100°C in Quartz cuvettes. The first derivative was used to determine the critical temperature (CT) (**see Table 1**) at which the **Q+Cu2+** and **Q+Fe2+** complexes became unstable in 2-Propanol at 50% and 80% concentrations (**Figure 2**).

**Table 1:** Critic Temperature of the Q+Cu2+ and Q+Fe2+ complexes.

| Complexes | CT (2-Prop50%) | CT (2-Prop80%) |
| --- | --- | --- |
| Q+Cu<sup>2+</sup> | 92.6°C | 87.6°C |
| Q+Fe<sup>2+</sup> | 91.6°C | 96.37°C |

**Figure 2:**
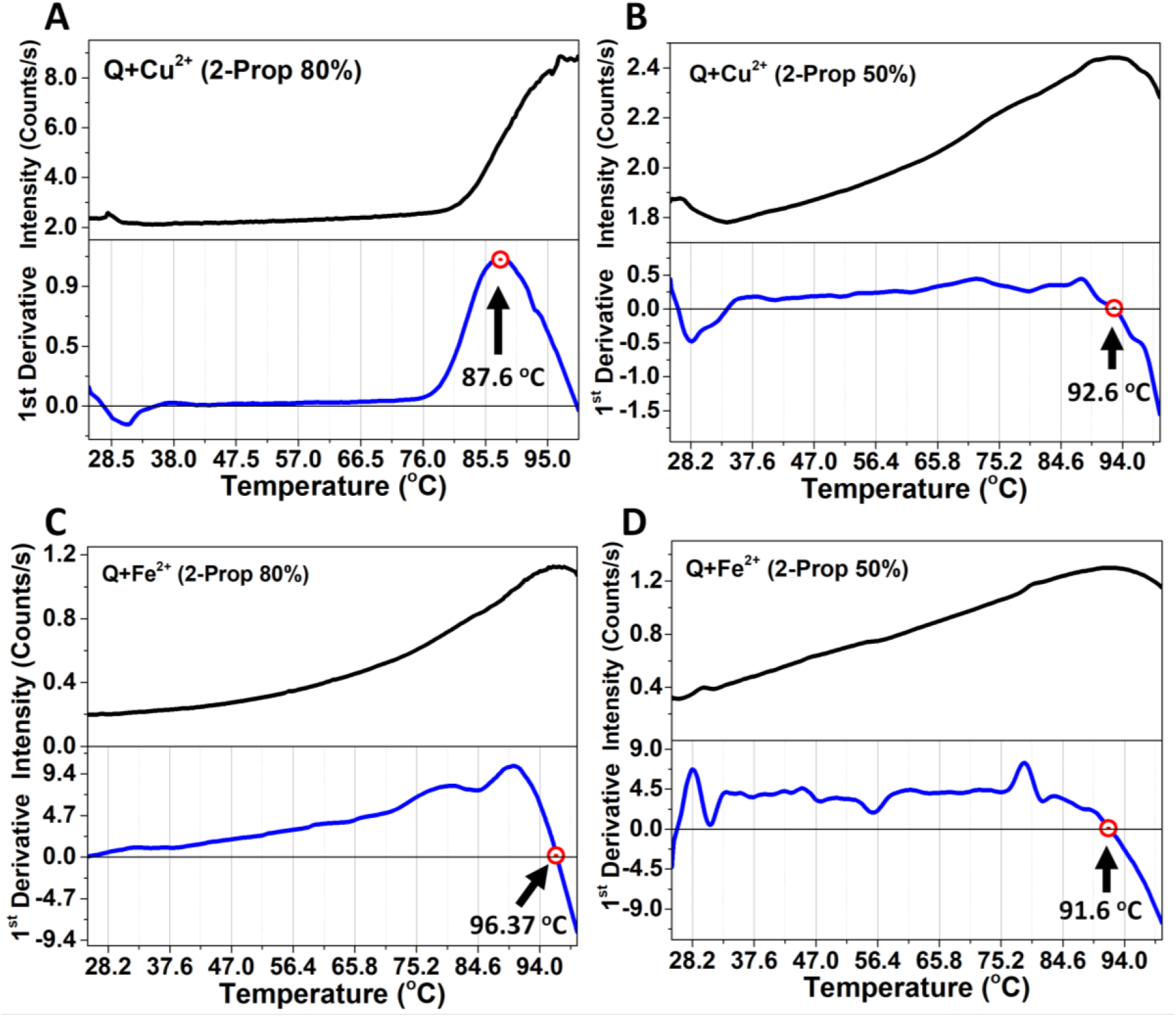
Determination of CT of Q/Cu^2+^/Fe^2+^ in 2-Propanol 80% and 2-Propanol 50%. The first derivative was used to determine the critical temperature (CT). Thermal Scan of **Q+Cu2+** in 2-Propanol 80% (CT = 87.6°C) (**A**) and 50% (CT = 92.6°C) (**B**). Thermal Scan of **Q+Fe2+** in 2-Propanol 80% (CT = 96.37°C) (**C**) and 50% (CT = 91.6°C) (**D**).

The results indicate that the stability is significantly affected by the solvent concentration. The case of **Q+Cu2+** dissolved in 2-Propanol 80% exhibited a lower critical temperature (CT = 87.6°C) compared to **Q+Cu**^**2+**^ in 2-Propanol 50% (CT = 92.6°C). This suggests that the complexation between **Q** and **Cu**^**2+**^ becomes unstable at a lower temperature in a more concentrated solvent (**Figure 2A**). In contrast, the stability of the quercetin-iron (**Q+Fe**^**2+**^) complex did not show a significant variation in the CT between the two 2-Propanol 80% and 2-Propanol 50%, which CT values were 96.37°C and 91.6°C respectively. This implies that the complexation of **Q** with **Fe**^**2+**^ is less influenced by solvent concentration as occurs with **Q** and **Cu**^**2+**^ (**Figure 2C and 2D**).

In **Table 1**, we summarize the values of the critic temperature from the complex in 2-Propanol 50% and 2-Propanol 80%.

### Temperature influence in the fluorescence properties of quercetin mixtures with Copper and Iron in 2-Propanol 50% and 2-Propanol 80%

In connection with the Thermal Scan (TS) (**see Figure 2**), we conducted fluorescence experiments to investigate the fluorescence behaviour of **Q** and its complexes with Copper (**Q+Cu**^**2+**^) and Iron (**Q+Fe**^**2+**^) at different temperature conditions, in two different solvent concentrations of 2-propanol (50% and 80%). The spectra were obtained under the next temperature conditions: 1-at 25°C before the thermal scan, 2-at 25°C after the thermal scan, and 3-at CT (Critic Temperature) (**see Table 1**), it is the temperature value where the complexes between **Q** and metal become unstable, and 4-at 25°C after acquiring the spectra at CT.

In the spectra of **Q** dissolved in both 2-propanol 50% and 80%, two prominent peaks were observed at 425 nm and 530 nm, corresponding to Ring A and Ring B of the quercetin molecule [31-33]. Compared with pure **Q**, these peaks exhibited variations in their intensity in the mixes **Q+Cu**^**2+**^ and **Q+Fe**^**2+**^ depending on the polarity of the solvent and influenced by the temperature conditions.

In the mix **Q+Cu**^**2+**^ dissolved in 2-Propanol 80%, the spectra taken at 25°C after the thermal scan as well as at CT = 87.6°C, showed a decreased intensity at 530 nm, while the peak at 425 nm completely disappeared in both measurements. However, at 25°C before the thermal scan, the peak at 425 nm reappeared with reduced intensity compared to pure **Q** (**Figure 3A**).

**Figure 3:**
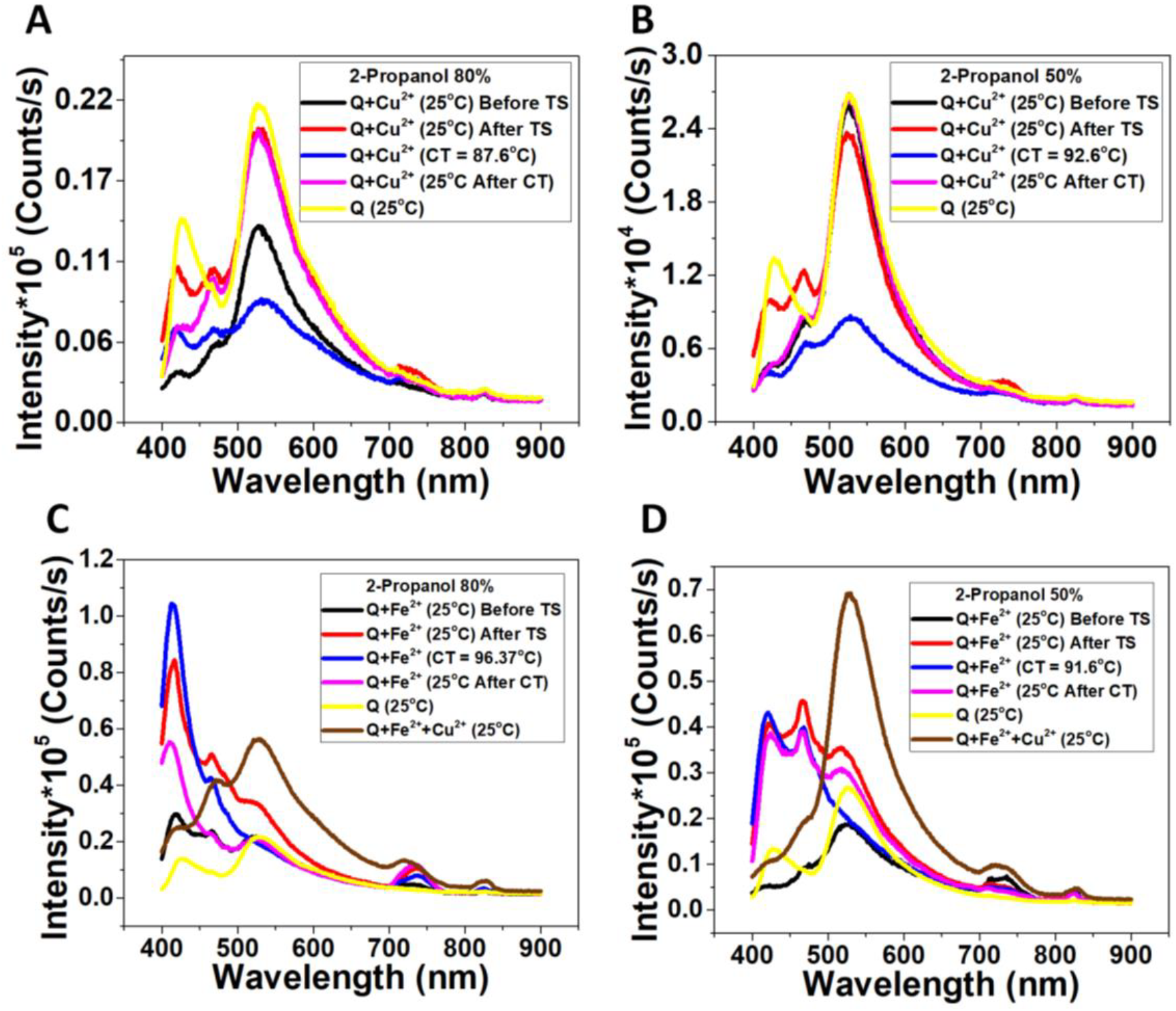
Influence of the temperature in the fluorescence properties of Q/Cu^2+^/Fe^2+^ in 2-Propanol 80% and 2-Propanol 50%. Similar to Q absorbance the fluorescence spectrum shows two peaks one corresponds to the Ring A at 425 nm and Ring B at 530 nm exciting at 370 nm. Fluorescence of Q compared with the fluorescence of **Q+Cu**^**2+**^ in 2-Propanol 80% (**A**) and 2-Propanol 50% (**B**). Fluorescence of Q compared with the fluorescence of **Q+Fe**^**2+**^ in 2-Propanol 80% (**C**) and 2-Propanol 50% (**D**).

In the case of **Q+Cu**^**2+**^ dissolved in 2-Propanol 50%, the spectrum taken at CT = 92.6°C similar to the analogue obtained in 2-Propanol 80% showed reduced intensity with respect to pure **Q**. However, at 25°C before the thermal scan, and 25°C after CT = 92.6°C the spectra exhibited a reduced intensity at 425nm, which differed from the spectrum at 25°C after the thermal scan where the peak at 425nm reappeared with reduced intensity (**Figure 3B**).

Surprisingly in the case of the **Q+Fe**^**2+**^ dissolved in 2-Propanol 80%, the peak at 530nm disappeared in all the spectra and the peak at 425nm notably increased the intensity compared to pure **Q**. Nevertheless, when **Q+Fe**^**2+**^ was dissolved in 2-Propanol 50% the peak at 530nm emerged from the spectrum at 25°C before the thermal scan (**Figure 3C**). This peak disappeared in the spectrum taken at CT = 91.6°C and reappeared as a shoulder in the spectra taken at 25°C after the thermal scan as well as the spectrum at 25°C after CT = 91.6°C. Peculiarly, after the thermal scan and the measurement at CT = 91.6°C a new peak was observed at 450nm which could reveal a new interaction between **Q** and **Fe**^**2+**^ in 2-Propanol 50% (**Figure 3D**).

In summary, our fluorescence results reveal distinctive changes in the intensity of peaks corresponding to Ring A and Ring B in **Q** and its complexes with **Cu**^**2+**^ and **Fe**^**2+**^, influenced by the solvent concentration and the critic temperature.

### Ratio scanning to explore interactions of Ring A and Ring B in the quercetin with Cu^2+^ and Fe^2+^

In connection with previous fluorescence emission results (**see Figure 3**), the Q exhibited characteristic peaks at 425 nm and 530 nm, corresponding to Ring A and Ring B in its structure. We investigated the interaction of **Q** with **Cu**^**2+**^ and **Fe**^**2+**^ by measuring the time-dependent fluorescence emission Ratio at those wavelengths (Ratio 425/530) in 2-Propanol 50% and 2-Propanol 80% at 25°C as well as at the critical temperature (CT) corresponding to each complexation occurred in 20 minutes (**see Figure 2 and Table 1**).

Our results show in both 2-Propanol 50% and 2-Propanol 80% at 25°C when the **Q** was in the presence of **Cu**^**2+**^ there was no significant increase in the Ratio 425/530 (**Figure 4C**). But at the CT = 92.6°C when the **Q+Cu**^**2+**^ was dissolved in 2-Propanol 50% there was no notable decrease in the Ratio 425/530 (**Figure 4A, blue plot**). However, at the CT = 87.6°C when the **Q+Cu**^**2+**^ was dissolved in 2-Propanol 80% a slight increase in the Ratio 425/530 was observed. This means that the interaction of the **Q** and **Cu**^**2+**^ may be partially influenced by the solvent concentration and the temperature (**Figure 4A, dark plot**).

**Figure 4:**
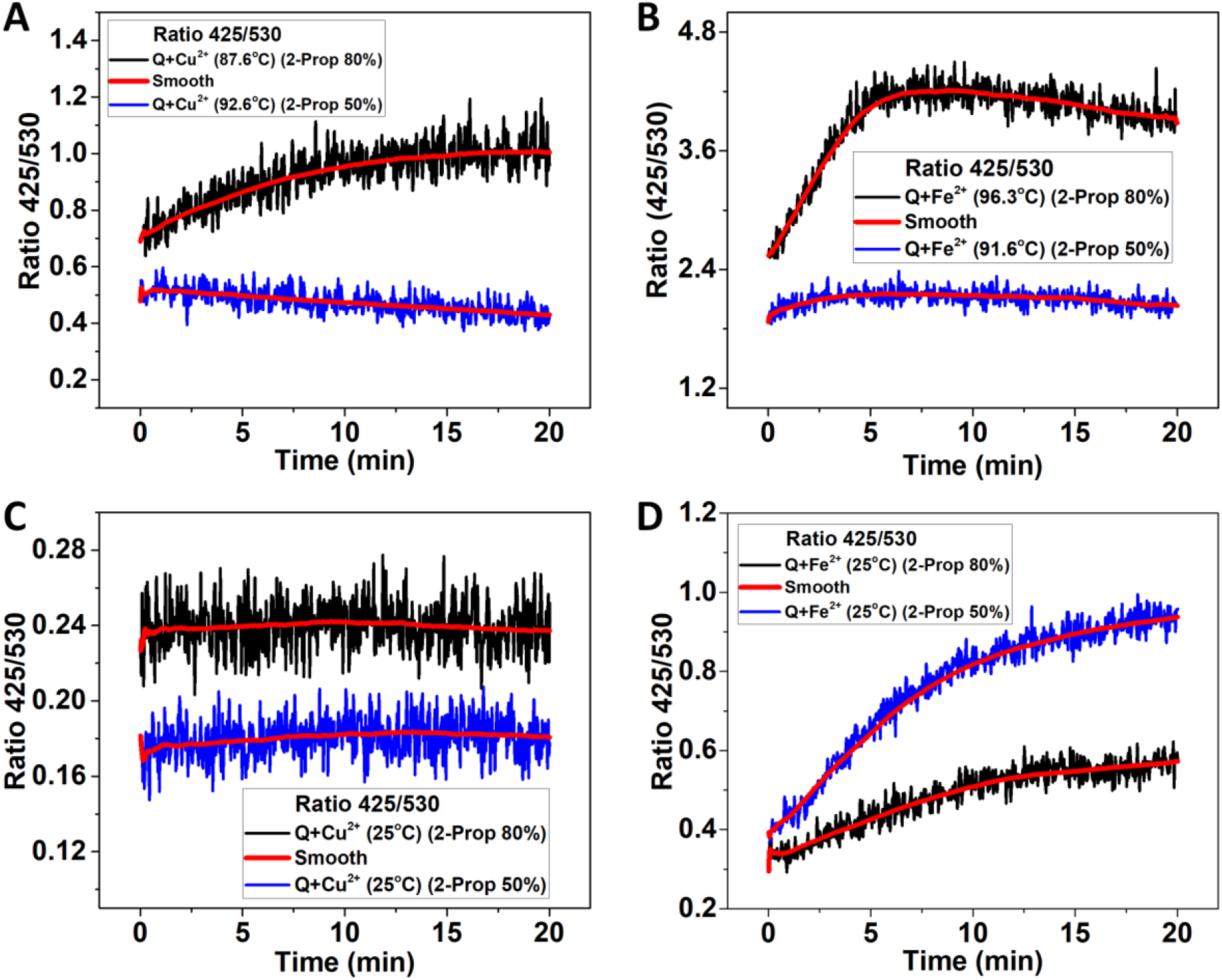
Exploring interactions of Ring A and Ring B in the quercetin with Cu^2+^ and Fe^2+^. Emission Ratio 425/530 of **Q+Cu**^**2+**^ at CT = 87.6°C in 2-Propanol 50% and CT = 92.6°C in 2-Propanol 80% (**A**) as well as 25°C (**C**). Emission Ratio 425/530 of **Q+Fe**^**2+**^ at CT = 91.6°C in 2-Propanol 50% and CT = 96.3°C in 2-Propanol 80% (**B**) as well as 25°C (**D**). In all cases, the Ratio 425/530 was measured during 20 minutes.

In the particular case of **Q+Fe**^**2+**^, after dissolving in both 2-Propanol 50% and 2-Propanol 80%, the Ratio 425/530 measured at 25°C increased ((**Figure 4D**). However, at CT = 91.6°C the Ratio 425/530 slightly decreased when quercetin was dissolved in 2-Propanol 50%, which suggests that the interaction behaviour at 25°C differs from CT = 91.6°C (**Figure 4B, blue plot**). Surprisingly, when **Q+Fe**^**2+**^ was dissolved in 2-Propanol 80% at the CT = 96.3°C, a strong increase in the Ratio 425/530 was observed in the first 5 minutes, and then a plateau behaviour appeared. This indicates a significant interaction which is stable during 15 min at this particular temperature (CT = 91.6°C).

### Erythema recovering after 24h by topically applying Q+Cu^**2+**^

The skin type classification was “Tanned” with an Individual Typology Angle (ITA) of **12.58**^**o**^, according to Bernerd et al. [34]. The solution **Q+Cu**^**2+**^ at 1μM:1μM ratio and dissolved in 2-Propanol 80% showed almost a constant scattering intensity during the thermal scanning from 25°C to 75°C (**see Figure 2A**). It is indicative that the interaction of **Q** with **Cu**^**2+**^ is more stable than the other solutions even in presence of **Fe**^**2+**^. The **Q** is known for its anti-inflammatory properties [35], additionally, **Cu**^**2+**^ have been suggested to participate in the melanin synthesis [15, 16]. In connection, three different areas on the skin were selected for induce the erythema and then evaluate the effect of **Q+Cu**^**2+**^ and pure **Q** by measure the erythema index reduction (**Figure 5B**).

**Figure 5:**
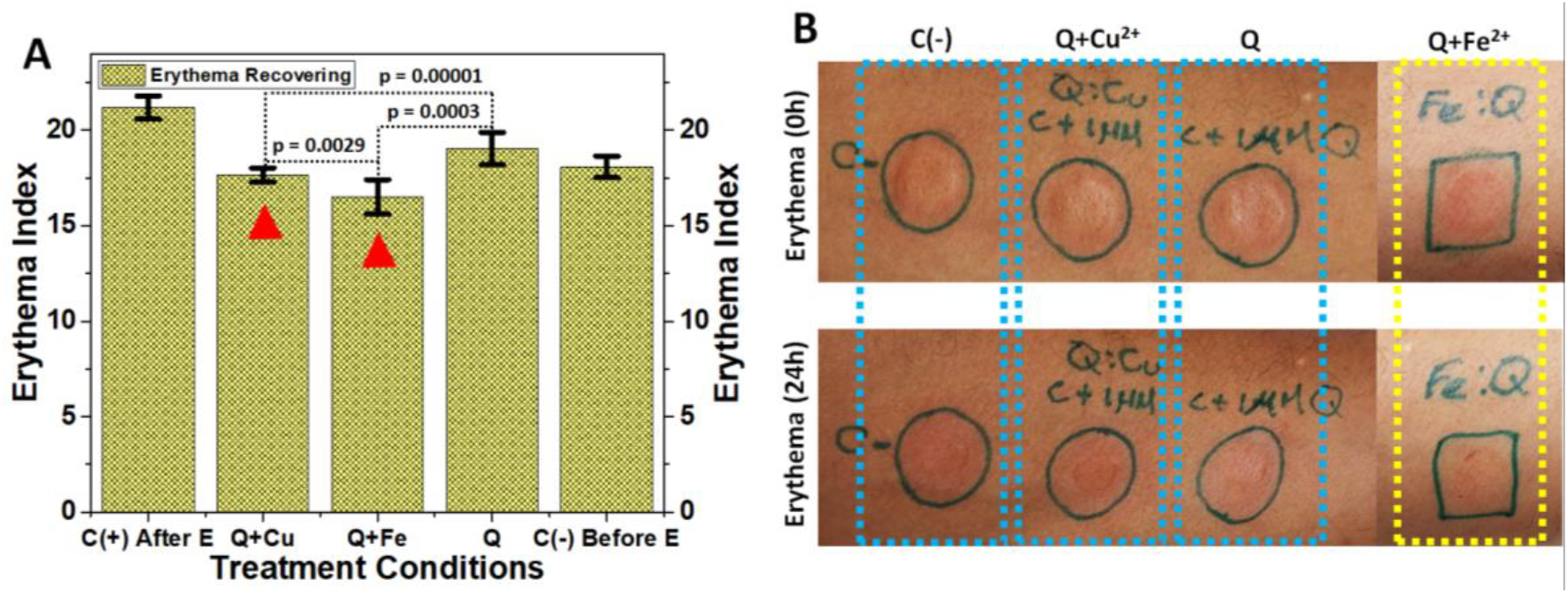
Erythema induction and recovery after 24h.The erythema index was measured using an electronic device, DSM III ColorMeter (Cortex Technology). **A:** Erythema index measured after treatment applied. The red arrows represent **Q+Cu**^**2+**^ and **Q+Fe**^**2+**^ treatments at (1μM:1μM) after 24h of erythema induction. The recovery is comparable with the negative control (**C(-)**) before erythema induction. **B:** Pictures of the skin application of the treatment at 0h (upper part) and 24h (lower part) of Erythema induction. Statgraphics program was used to statistical analysis.

Thus, two negative controls (2-Propanol 80%) were used before and after erythema induction (**Figure 5A**).

After 24h, the erythema index of each experimental area was measured using an electronic device. The t-student statistical analysis was using Statgraphics. According to this analysis, the erythema index showed a significant reduction in both **Q+Fe**^**2+**^ and **Q+Cu**^**2+**^ treatments respect to pure **Q** treatment area with p-values of 0.0003 and 0.00001 respectively. However, significant difference also was observed between **Q+Fe**^**2+**^ and **Q+Cu**^**2+**^. This suggests that the combination of **Q** with **Fe**^**2+**^ and **Cu**^**2+**^ separately has a synergic effect in the erythema reduction.

## DISCUSSION

The spectroscopic results presented in this study provide new insights into the interactions between quercetin and two biocompatible metal ions (**Cu**^**2+**^ and **Fe**^**2+**^). These interactions are of interest because of the implications of **Q** in the reduction of inflammation in the context of skin health [10, 11]. In connection with this, **Cu**^**2+**^ ion skin effects have been studied more than **Fe**^**2+**^. **Cu**^**2+**^ has a clear physiological role in the skin [4], highlighting dermal fibroblast proliferation [12], as a cofactor of skin enzymes such as superoxide dismutase, which protects against free radicals [13, 14] as well as tyrosinase and melanin enzymatic biosynthesis, with a central role in skin pigmentation [15, 16]. On the other hand, **Fe**^**2+**^ ions play an important role in collagen metabolism as well as procollagen-proline dioxygenase [17], and their anti-inflammatory effects on the skin are not as clear as those of iron chelators that do not include Q [18, 36]. However, recent studies have reported the anti-inflammatory effect of **Q+Fe**^**2+**^ nanoparticles in swollen joints of mouse models of rheumatoid arthritis [21]. Herein, we have addressed the application to reduce the induced skin erythema by using both **Q+Cu**^**2+**^ and **Q+Fe**^**2+**^ (**See Figure 5**).

For the purposes of this study, we found that in both 2-Propanol 80% and 50% provide good solubility for both **Q+Cu**^**2+**^ and **Q+Fe**^**2+**^ at a studied proportion of 1:1. Additionally, this allowed us to evaluate the effect of solvent polarity on the spectroscopic properties of the **Q/Cu**^**2+**^**/Fe**^**2+**^ mixtures. By using this proportion (1:1), we can determine which ions are more suitable for interacting with a specific part of the **Q** molecule, Ring A or Ring B. It is important to provide a better understanding of which part of the **Q** molecule is more important for triggering the anti-inflammatory effect on the skin under the tested experimental conditions.

However, with respect to the usability of 2-propanol in skin treatments, it is important to highlight that the concentration is important. Thus, lower concentrations of approximately 70% are used in healthcare because they show less drying and irritant effects on the skin [37, 38] owing to a mechanism that affects the stiffness of collagen [39]. In addition, topical application for preoperative skin studies demonstrated poor penetration of 2-propanol 70% into the deeper layers of the skin, and non-skin irritation was reported [40, 41]. Thus, the percentage of the experimental solution used in this study (2-Propanol 80%) to test the skin erythema reduction may be used as a suitable negative control (**See Figure 5**).

Respect to **Q+Cu**^**2+**^ interaction, there is a significant change in the fluorescence emission spectrum at 530 nm compared to **Q+Fe**^**2+**^ and depending on the 2-Propanol concentration (Figure 3A, B, C and D, Black color). This increased emission intensity at 530 nm suggest a coordination of **Cu**^**2+**^ with the carbonyl oxygen atom, -OH3 and -OH5 group in the **Q** molecule, Ring A-C (Benzoyl group) (Figure 3A and B, Black color), which match with the reported data [22, 23, 42]. Thus, according to Joshi et al. [43] the -H3’ and -4’OH in Ring B (cinnamoyl group) of the **Q** molecule may form a complex with COX-2, interacting through hydrogen bonds with Ser530 and Tyr385, respectively. The COX-2 (cyclooxygenase 2) is an enzyme with a central role in the inflammatory pathway [35, 44]. In addition, **Q** can reduce the activation of NF-kB by blocking UV irradiation during topical applications [45, 46]. In agreement with the anti-inflammatory properties of **Q** [35, 47], our results suggest that both metal ions, **Cu**^**2+**^ and **Fe**^**2+**^, interact with the **Q** molecule, which improves the inhibition effect on COX-2, enhancing the anti-inflammatory properties of **Q**. However, statistical analysis showed that **Q+Fe**^**2+**^ chelate had a better anti-inflammatory effect than **Q+Cu**^**2+**^ and pure **Q** (see Figure 5A). This result is interesting because the **Q+Fe**^**2+**^ interaction showed a contrasting behavior in the fluorescence emission spectra between 2-Propanol 80% and 50% in computation with similar solutions in the **Q+Cu**^**2+**^ chelate (see Figure 3C and 3D). This behavior is consistent with that reported by Satterfield et al. [26], as well as the strong affinity of **Fe**^**2+**^ for **Q** molecules, which is visible in a ternary mixture where the predominant color is brown (see Figure S1). Thus, our results suggest a preference for the coordination of Fe2+ with the carbonyl oxygen atom, -OH3, and -OH5 groups in the **Q** molecule, Ring A-C (benzoyl group) dissolved in 2-Propanol 50% (see Figure 3D, black color). However, when **Q** is dissolved in 2-Propanol 80%), our fluorescence emission results suggest that coordination occurs predominantly for -OH4’ and -OH3’ in the cinnamoyl group (Ring B) of the **Q** molecule according to reported data [27-29]. Although, the fluorescence spectrum shows a reduction in the emission at both wavelength (425 nm and 530 nm) compared with the spectrum obtained in 2-Propanol 50% (see Figure 3C and 3D Black color). The differential fluorescence emission behavior depends on two factors: 1) the solvent concentration influences the pH of the solution, favoring the formation of a reversible p-quinone methide structure between the carbonyl oxygen atom, -OH3, or -OH5 group in the **Q** molecule, which is in agreement with previously reported data [48-52]. 2-The high reductive decomplexation capacity of **Fe**^**2+**^ over chelated **Cu**^**2+**^ allows it to form chelates more easily than **Cu**^**2+**^ [30], which supports our observations of the differential fluorescence emission of **Q+Fe**^**2+**^ in 2-Propanol 50% and 2-Propanol 80%) (see Figure 3C and 3D black color). In addition, the improved anti-inflammatory effect of the **Q+Fe**^**2+**^ chelate responds to the role of the hydroxyl groups stabilizing the interaction of **Q** with **Fe**^**2+**^, highlighting that the **Fe**^**2+**^ ion is required for the enhanced anti-inflammatory effects of **Q** (**see Figure 5A**).

## CONCLUSION

In summary, the 2-propanol concentration influenced the spectroscopic properties when the Q molecule chelated biocompatible metal ions (**Fe**^**2+**^ and **Cu**^**2+**^). However, two well distinguishable fluorescence emission peaks (425 nm and 530 nm) emerged in 2-propanol 80% depending what metal ion is involved in the chelate. This spectroscopic behavior reveals the specific part of the Q molecule implicated in the interaction with **Cu**^**2+**^ and **Fe**^**2+**^, as well as the strength of the interaction between them, with **Q+Fe**^**2+**^ + being the most stable chelate compared with **Q+Cu**^**2+**^. Our results suggest that both experimental conditions (**Q+Fe**^**2+**^ and **Q+Cu**^**2+**^) with clear anti-inflammatory properties effectively enhanced the individual anti-inflammatory properties of Q molecules in induced skin erythema (skin type, Tanned). However, the effect of **Q+Fe**^**2+**^ chelate was statistically greater than the **Q+Cu**^**2+**^ chelate. In addition, we propose that the number of hydroxyl groups involved in chelating formation plays a pivotal role in mediating the interaction with metal ions to trigger the anti-inflammatory effect of the chelates.

## Abbreviations and Symbols

Fe^2+^: Q (Quercetin),
Cu^2+^: (Iron II),
SLS (Static Light Scattering): (Copper II)

---

**Figure S1:**
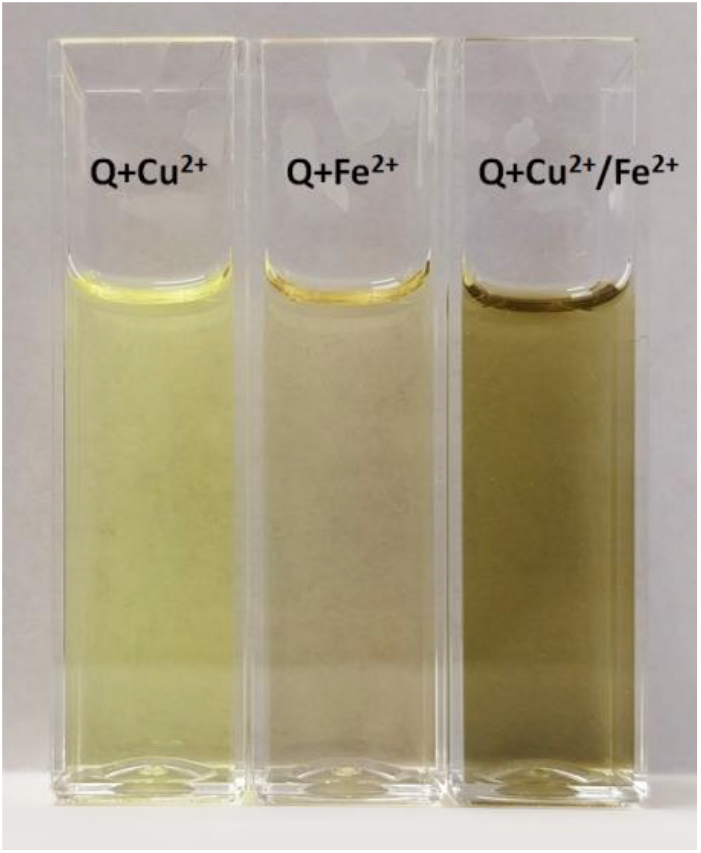
Predominant color in the Q+Fe^2+^, Q+Cu^2+^, Q+Cu^2+^/Fe^2+^.

